# Synthetic Lethal Interactions Prediction Based on Multiple Similarity Measures Fusion

**DOI:** 10.1101/2020.08.03.235366

**Authors:** Yu-Qi Wen, Lian-Lian Wu, Xiao-Xi Yang, Bo-Wei Yan, Song He, Xiao-Chen Bo

## Abstract

The synthetic lethality (SL) relationship arises when a combination of deficiencies in two genes leads to cell death, whereas a deficiency in either one of the two genes does not. The survival of the mutant tumor cells depends on the SL partner genes of the mutant gene, so the cancer cells could be selectively killed by inhibiting the SL partners of the oncogenic genes but normal cells not. Therefore, developing SL pairs identification methods is increasingly needed for cancer targeted therapy. In this paper, we proposed a new approach based on similarity fusion to predict SL pairs. Multiple types of gene similarity measures are integrated and k-NN algorithm are applied to achieve the similarity-based classification task between gene pairs. As a similarity-based method, our method demonstrated excellent performance in multiple experiments. Besides the effectiveness of our method, the ease of use and expansibility can also make our method more widely used in practice.

## 1. Introduction

The synthetic lethality (SL) relationship occurs in two genes when the perturbation of two genes lead to cell death or a sharp decline in cell viability ^[1,2]^. This form of cell killing is based on the interactions of two genes. The cell is viable when either gene is mutated alone, but is lethal when the combination of mutations occurs in both genes^[3]^. The survival of mutant tumor cells depends on the SL partner genes of the mutant gene, so SL partners can potentially serve as drug targets when the driver genes or oncogene cannot be targeted^[4]^. The cancer cells could be selectively killed by the chemical inhibition of the SL partners of the oncogenic genes but normal cells not^[5]^. For instance, BRCA1- and BRCA2-deficient cells are sensitive to treatment with inhibitors of poly (ADP-ribose) polymerase (PARP)^[6,7]^. This SL pair is used to treat breast and ovarian cancers where BRCA1 or BRCA2 are mutated^[8,9]^. Therefore, identifying clinically available SL pairs for screening potential targets is important to improve the efficacy of anticancer treatment^[4,10]^.

High-throughput screening (HTS) technology has been developed to identify potential SL pairs, such as chemical screening^[11]^, pooled siRNA or shRNA screening^[12,13]^ and the CRISPR/Cas9 system for gene knockdown^[14,15]^, producing large number of available SL data in reasonable time and at low costs^[16,17]^. Recently, a comprehensive database, SynLethDB with 34,089 SL pairs of different species was performed, which collected SL pairs from HTS experiments, experimental literatures and so on^[18]^. Although such screening techniques are effective approaches, testing the complete SL space with HTS is unfeasible.

Computational approaches have been developed to offer the possibility to efficiently explore the large SL space. The available HTS SL data can be leveraged to generate accurate predictive models. The *in vitro* and *in vivo* research can be guided by the reliable predictions. To predict potential SL pairs, many computational methods perform individual analysis of data sources from evolutionary characteristics^[19,20]^, transcriptomic profiles^[21]^, interaction network^[22]^, to cancer patient data^[10,23]^. Moreover, recent development of neural network has also attracted the attention of researchers. Wan *et al*. proposed a machine learning framework for cell-line-specific synthetic lethality prediction, which was a semi-supervised neural network-based method called EXP2SL to identify SL interactions from the L1000 gene expression profiles^[24]^. Compared with the above methods that use only a single data type, integrating multiple data types enables more informative and comprehensive analysis of SL interactions, and multiple sources of evidence pointing to the same result are less likely to lead to false positives. Therefore, methods for integrating multi-dimensional SL gene related data are increasingly needed. Liany *et al*. applied collective matrix factorization to integrate multiple heterogeneous data, which in turn were used for prediction SL interactions through matrix completion^[25]^. Although it is powerful, the method that operates with high-dimensional feature × sample matrices have scalability drawbacks, making it sensitive to feature dimension. When the feature dimension is too large, the computational complexity will increase. And feature preselection step may also affect model performance.

Instead of processing largescale matrices constructed over a large number of features, similarity-based integration strategy uses similarity matrix as a basis and is not sensitive to feature dimension and preselection. In this work, we proposed a new approach for predicting SL gene pairs, through integrating the similarity measures based on the gene expression profile, protein sequence, protein–protein interaction (PPI) network, co-pathway and Gene Ontology (GO). We applied the k-NN algorithm to achieve the similarity-based classification task between gene pairs. Our approach was trained on the SynLethDB, which is a large publicly SL database. Next, we compared the performance of our model with the model based on each single similarity measure. Additionally, to benchmark the performance of our approach, we compare the results to Probability Ensemble Approach (PEA) algorithm, another similarity-based algorithm which get great performance in the classification task^[26]^. Overall, we found that our approach with the integrated similarity measure can predict SL gene pairs with higher performance of an AUROC of 0.85 compared to other methods. Next, we applied our approach to predict novel SL gene pairs. We found that the RAS genes (i.e. KRAS, NRAS, HRAS) have the largest number of SL partners both in the training set and the predicted top 3,000 SL pairs. We further employed a pathway enrichment analysis and calculated the ATC distribution of drugs for the RAS genes’ SL partners. The results show that these partner genes might be promising targets to achieve synthetic lethality for the cancer cells in the targeted therapy.

## 2. Materials and methods

### 2.1 Dataset

#### 2.1.1 Synthetic lethality (SL) data

To collect known SL information, we used SynLethDB, a public database for SL interactions^[18]^. SynLethDB (http://histone.sce.ntu.edu.sg/SynLethDB/) contains 19,952 human SL pairs collected from biochemical assays, other related databases, computational predictions by DAISY^[23]^ and text mining results.

For the negative samples (i.e., pairs that are not SL), we first extracted the PPI subnetwork from the complete PPI network^[27]^ based on genes involved in SL interaction in SynLethDB. Then we excluded SL interaction in SynLethDB from this PPI subnetwork, and we constructed our negative sample set by randomly selecting pairs from the remaining interaction. We make sure that the negative training set and the positive training set have the same number of samples.

#### 2.1.2 Feature data for gene-gene similarity measures

To calculate gene-gene similarity measures, we collected feature data from multiple data sources. We extracted gene expression profile data from the Library of Integrated Network-Based Cellular Signatures (LINCS) project, which is a mutual fund project administered by the National Institutes of Health (NIH). This project uses L1000 technology to generate approximately one million gene expression profiles^[28]^. In this study, we used the gene knockdown transcriptome data in the database.

The protein sequence data were extracted from the UniProt database, which provides high-quality, freely accessible protein sequence data^[29]^.

For PPI network data, we used the PPI network provided by the article^[27]^. It integrates PPI networks from 7 sources, including a total of 141,296 associations between 13,460 proteins.

The gene-pathway association data is collected from the Comparative Toxicogenomics Database (CTD) database^[30]^. In all pathway data, we only use the Reactome pathway data to make sure that each gene has a fixed-length pathway vector.

### 2.2 Model construction

The overall framework is illustrated in Fig.1. It can predict whether a pair of genes is SL pair by integrating multiple properties of genes. The basic hypothesis of our approach is that the more similar gene pairs are, the more likely they are to have the same SL characteristics. The framework consists of four parts: (1) First, given a pair of query genes, seven gene-based similarity measures of the two genes to the known SL pairs are calculated. (2) Then, the SNF algorithm is applied to fuse the seven types of similarity measures into one integrated similarity measure. (3) In the third part, the similarity measure for gene pairs is defined based on gene-gene similarity measures. Therefore, the similarity measures between the query gene pair and the known SL gene pairs is calculated. (4) After that, k-NN algorithm is performed to determine whether the query gene pair is SL pair based on the training set.

**Fig.1.**
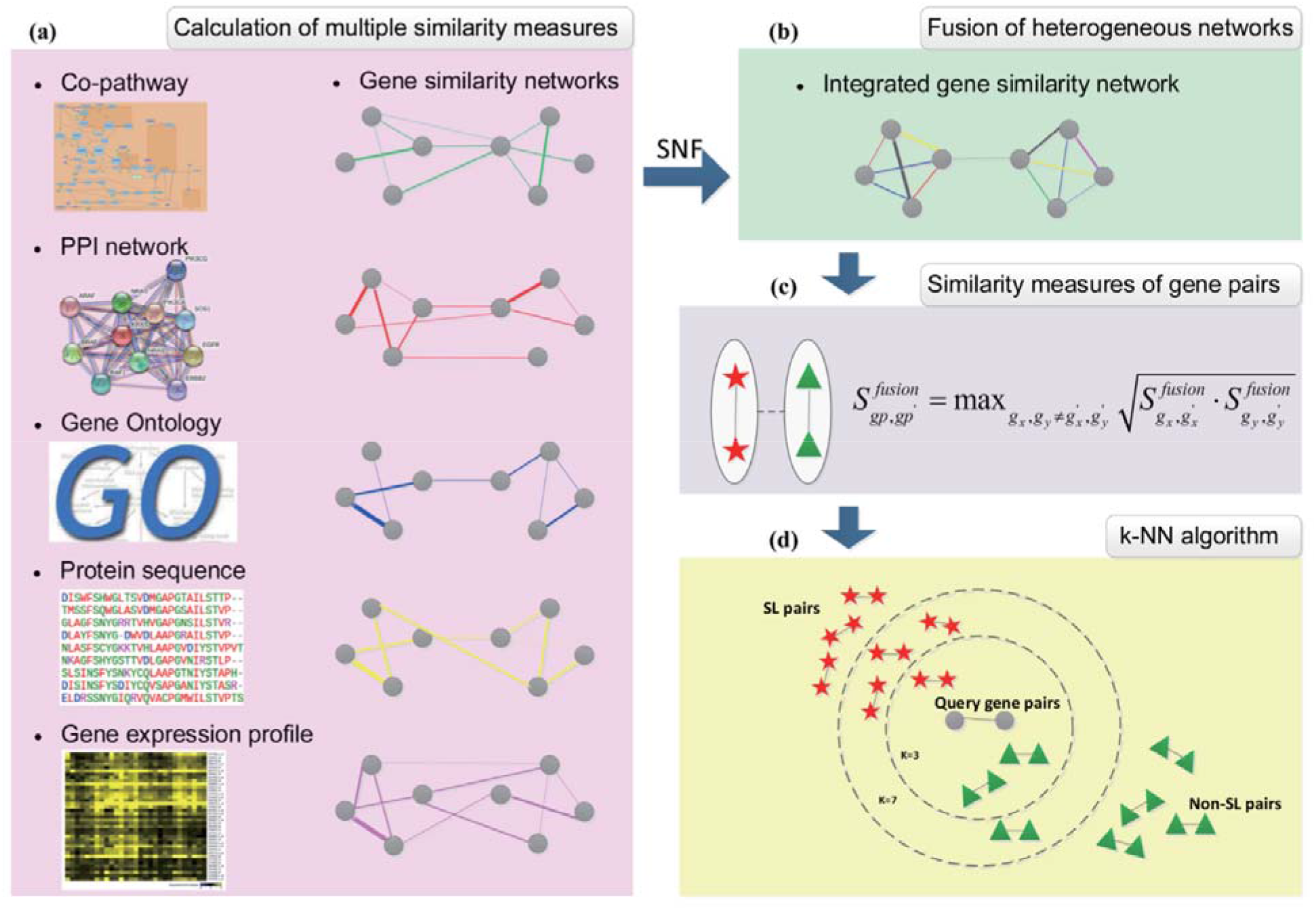
The framework of our approach consists of four parts: (a) calculating the seven gene-based similarity measures (three types of GO-based) of the query genes to the known SL genes. (b) Fusing the seven types of similarity measures into one integrated similarity measure with the SNF algorithm. (c) Calculating the similarity measures between the query gene pair and the known SL gene pairs based on the integrated similarity measures. (d) Applying the k-NN algorithm to determine whether the query gene pair is SL pair based on the training set.

#### 2.2.1 Gene-gene similarity measures

We defined and computed seven gene–gene similarity measures including the similarities based on gene expression profile, gene encoded protein sequence, PPI network, co-pathway, Gene Ontology Biological Process (GOBP), Gene Ontology Cellular Component (GOCC) and Gene Ontology Molecular Function (GOMF). All similarity measures were normalized to be in the range [0, 1].

##### Similarities based on gene expression profile

We used the method named Gene Set Enrichment Analysis (GSEA) as a similarity measure^[32]^.

The specific calculation process of this method is as follows: first of all, we constructed Prototype Ranked List (PRL) of genes for each gene knockdown sample in the dataset. The PRL is a list of genes ranked according to their differential expression following gene knockdown treatment, from the most up-regulated to the most down-regulated^[31]^. Then we selected the top-ranked 250 genes and the bottom-ranked 250 ones (denoted by *p* and *q* respectively) from each PRL to form the signature for each gene knockdown sample. For gene *x* and gene *y*, the enrichment score of the gene *x* signature {*p*, *q*}, with respect to the PRL of gene *y*, is defined as:

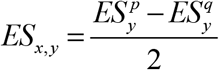

Similarly, we can get *ES*_*y,x*_, the enrichment score of the gene *y* signature, with respect to the PRL of gene *x*. At last, we get the similarity measure between *g*_*x*_ and *g*_*y*_ based on gene expression profile:

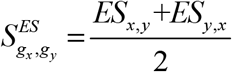

##### Similarities based on gene encoded protein sequence

We calculated the similarity measure between pairwise proteins sequence encoded by *g*_*x*_ and *g*_*y*_ using SW algorithm (Smith-Waterman algorithm):

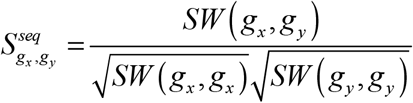

*g*_*x*_ represents the protein sequence encoded by gene *x*, *g*_*y*_ represents the protein sequence encoded by gene *y*. *SW*(*g*_*x*_, *g*_*y*_) is the Smith-Waterman sequence alignment score of protein sequence^[33]^.

##### Similarities based on protein–protein interaction (PPI) network

We used the following formula to calculate the similarity measure between gene *g*_*x*_ and gene *g*_*y*_ :

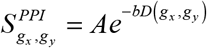

where, *D*(*g*_*x*_, *g*_*y*_) represents the shortest distance between *g*_*x*_ and *g*_*y*_ on the PPI network. We used Dijkstra’s algorithm to calculate the shortest distance in the network. According to Perlamn *et al.* study, we set *A* = 0.9 × *e*, *b* = 1 ^[34]^.

##### Similarities based on co-pathway

We obtained the association data between genes and 1,860 Reactome pathways from the CTD database^[30]^. Each gene has an 1,860-dimensional feature vector, represented by *P*(*g*). We used the Tanimoto coefficient to calculate the similarity measure between gene *g*_*x*_ and gene *g*_*y*_ :

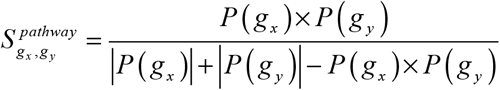

where, |*P*(*g*_*x*_)| and |*P*(*g*_*y*_)| represent the number of pathways in which gene *g*_*x*_ and gene *g*_*y*_ respectively. *P*(*g*_*x*_) × *P*(*g*_*y*_) represents the number of involved pathways shared by gene *g*_*x*_ and gene *g*_*y*_.

##### Similarities based on Gene Ontology (GO)

We used the R package GOsemsim^[35]^ to calculate the GO semantic similarity of molecular function (MF), biological process (BP), and cellular component (CC).

Given two sets of GO terms that annotate genes *g*_*x*_ and *g*_*y*_, *GO*_*x*_ = {*go*_*x*1_, *go*_*x*2_, …, *go*_*xm*_} and *GO*_*y*_ = {*go*_*y*1_, *go*_*y*2_, …, *go*_*yn*_}, the similarity measure of *g*_*x*_ and *g*_*y*_ can be calculated as follows:

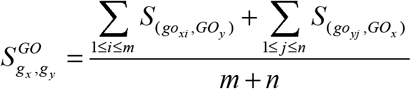

where, *S*_(*go*,*GO*)_ is calculated as follows:

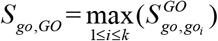

It represents the maximum semantic similarity between term *go* and any of the terms in set *GO*. For the semantic similarity 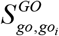, we used the method proposed by Wang *et al.*, a graph-based method to compute semantic similarity using the topology of the GO graph structure^[36]^.

#### 2.2.2 Fusion of heterogeneous features

We used the R package SNFtool to achieve SNF, a useful computational method for data integration in the field of disease subtype identification^[37]^. It can make full use of common and complementary information across data types by integrating data in a non-linear way.

In our model, by calculating the above similarity measures, we can obtain seven similarity matrixes *S*_1_, *S*_2_, … *S*_7_. Then the seven similarity matrixes can be normalized as follows:

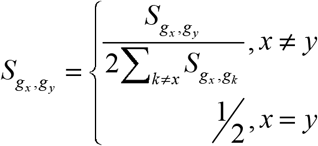

*g*_*x*_ and *g*_*y*_ represent gene samples. Moreover, for each similarity matrix, a local affinity matrix A is defined:

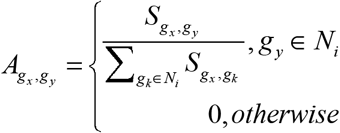

*N*_*i*_ denotes a set of neighbors of *g*_*x*_ including *g*_*x*_ in a similarity network.

We integrated the multiple gene similarity networks with SNF, which iteratively updated each of seven similarity matrixes as follows:

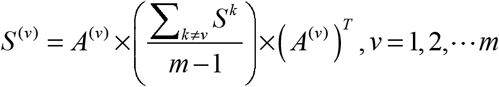

Here, *n* = 7, because we have seven types of data. For example, let 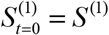, *P*_*t+1*_ is updated as 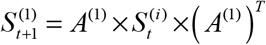,*i* = 2,3,…7 .

After *t* steps, the final gene similarity matrix is computed as:

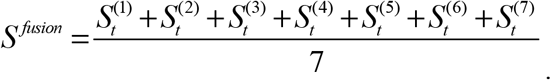

#### 2.2.3 Similarity measures for gene pairs

After obtaining the integrated gene-gene similarity matrix, the similarity measure between a query gene pair (*g*_*x*_, *g*_*y*_) and a known gene pair 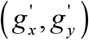 is defined as follows:

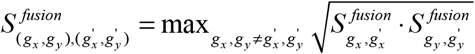

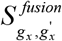 and 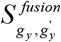 (and symmetrically 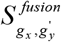 and 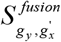) donate the integrated gene-gene similarity measures. Then, these two similarity measures are combined by calculating their geometric mean^[34]^. By taking the larger of 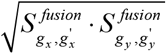 and 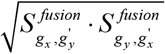, we get the final similarity measure between a query gene pair (*g*_*x*_, *g*_*y*_) and a known gene pair 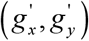, 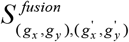.

#### 2.2.4 k-NN algorithm

K-Nearest Neighbors algorithm (k-NN) is a non-parametric method used for classification^[38]^. In k-NN classification, a sample is classified by a plurality vote of its neighbors.

For SL prediction, we first obtained the k nearest neighbors of the query gene pair by similarity ranking between the query gene pair and the known SL gene pairs. Then the query gene pair is assigned to the class (SL or non-SL) most common among these k nearest neighbors.

For k-NN, we determined the best k by cross-validation. By sampling across the range of *k* choices, we set *k* = 19 which led to the highest AUROC in cross-validation.

### 2.3 Experiment

To evaluate the predictive performance of our model, we used a 10-fold cross validation. We compared the model performance before and after SNF. And we compared the performance of our model with the PEA (Probability Ensemble Approach) algorithm, which is a probability-based model for classification^[26]^. PEA combines multiple similarity measures into one score through Bayesian network. By putting the integrated score into a random score distribution, the score can be converted to a *P* value (ranging from 0 to 1). This resulting *P* value represents the probability of a given score that better to be observed from random data the final classification is determined by comparing the *P* value with a preset threshold. We choose PEA as our benchmark because it also inferred association based on similarity.

## 3. Results

### 3.1 Performance evaluation

To find data-driven motivation and verify the basic hypothesis for our approach, we further check the statistical distribution of these similarity measures of SL and non-SL gene pairs in the training set (Fig. 2). The difference is observed in the fusion similarity feature between SL and non-SL gene pairs, which is one of the reasons for the good classification effect in our approach.

**Fig.2.**
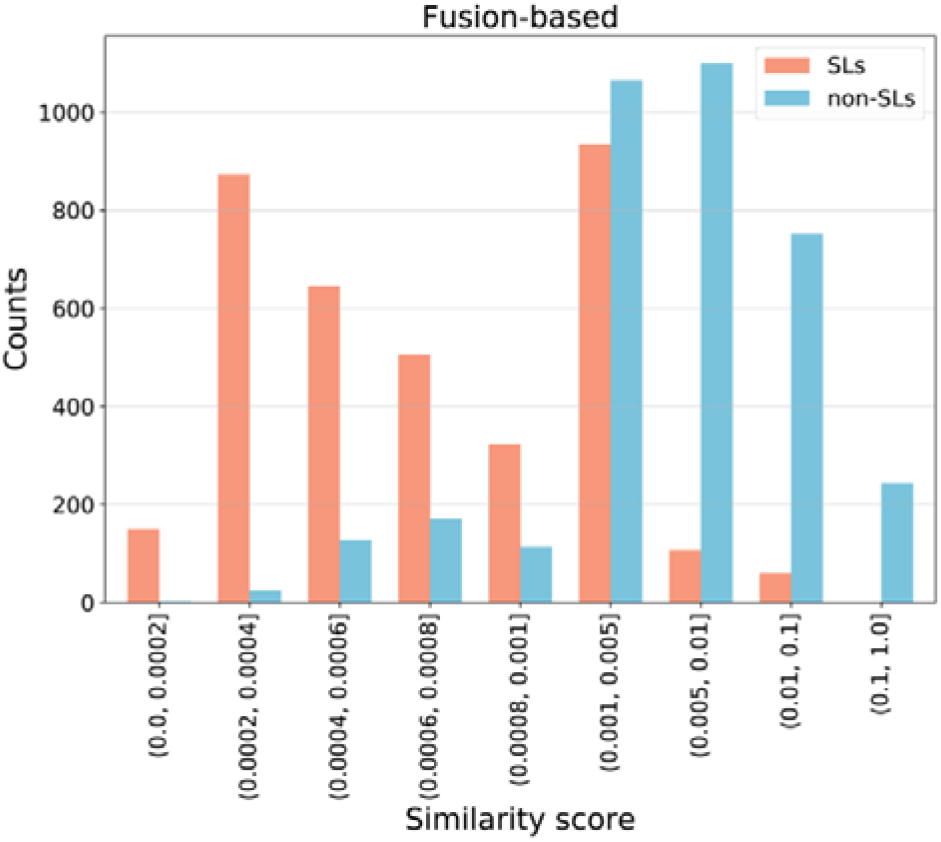
The statistical distribution of the fusion of seven similarity measures.

To quantitatively assess the performances of our model with the integration of all seven similarity measures or each single similarity measure in predicting SL pairs, we performed a 10-fold cross validation accompanied with the receiver operating characteristic (ROC) and the precision recall (PR) curves analysis in the SL data (Fig.3(a)~3(b)). Moreover, to characterize the predictive performance of our approach and give comparable measures, we also used the performance measures that are typical for classification tasks: area under the receiver operator characteristics curve (AUROC), area under the precision recall curve (AUPR), accuracy, precision, recall and F1 score (Table 1). After the selection of parameters for the k-NN classifier, our model with the integrated similarity measure achieved test AUROC of 0.85 and AUPR of 0.86, which exhibited better performance than those with single similarity measure. Furthermore, among all other evaluation metrics including accuracy, precision, recall and f1 score, our method shows better performance than any other single similarity measure-based model (Table1).

**Table 1.** Performance comparison of the k-NN and PEA algorithm based on the integrated similarity measures and seven single similarity measures for predicting SL gene pairs.

| Method | AUROC | AUPR | Precision | Recall | F1 | Accuracy |
| --- | --- | --- | --- | --- | --- | --- |
| **Fusion-based k-NN** | **0.848** | **0.861** | **0.825** | 0.670 | **0.739** | **0.764** |
| GOBP-based k-NN | 0.745 | 0.754 | 0.708 | 0.607 | 0.653 | 0.678 |
| GOCC-based k-NN | 0.663 | 0.623 | 0.708 | 0.428 | 0.533 | 0.626 |
| GOMF-based k-NN | 0.531 | 0.531 | 0.602 | 0.121 | 0.201 | 0.520 |
| Co-pathway-based k-NN | 0.761 | 0.779 | 0.708 | 0.659 | 0.682 | 0.693 |
| PPI-based k-NN | 0.500 | 0.472 | 0.542 | 0.812 | 0.650 | 0.562 |
| Gene expression-based k-NN | 0.580 | 0.589 | 0.528 | 0.716 | 0.607 | 0.537 |
| Protein sequence-based k-NN | 0.762 | 0.791 | 0.726 | 0.609 | 0.662 | 0.689 |
| Fusion-based PEA | 0.723 | 0.733 | 0.585 | **0.914** | 0.713 | 0.633 |

**Fig.3.**
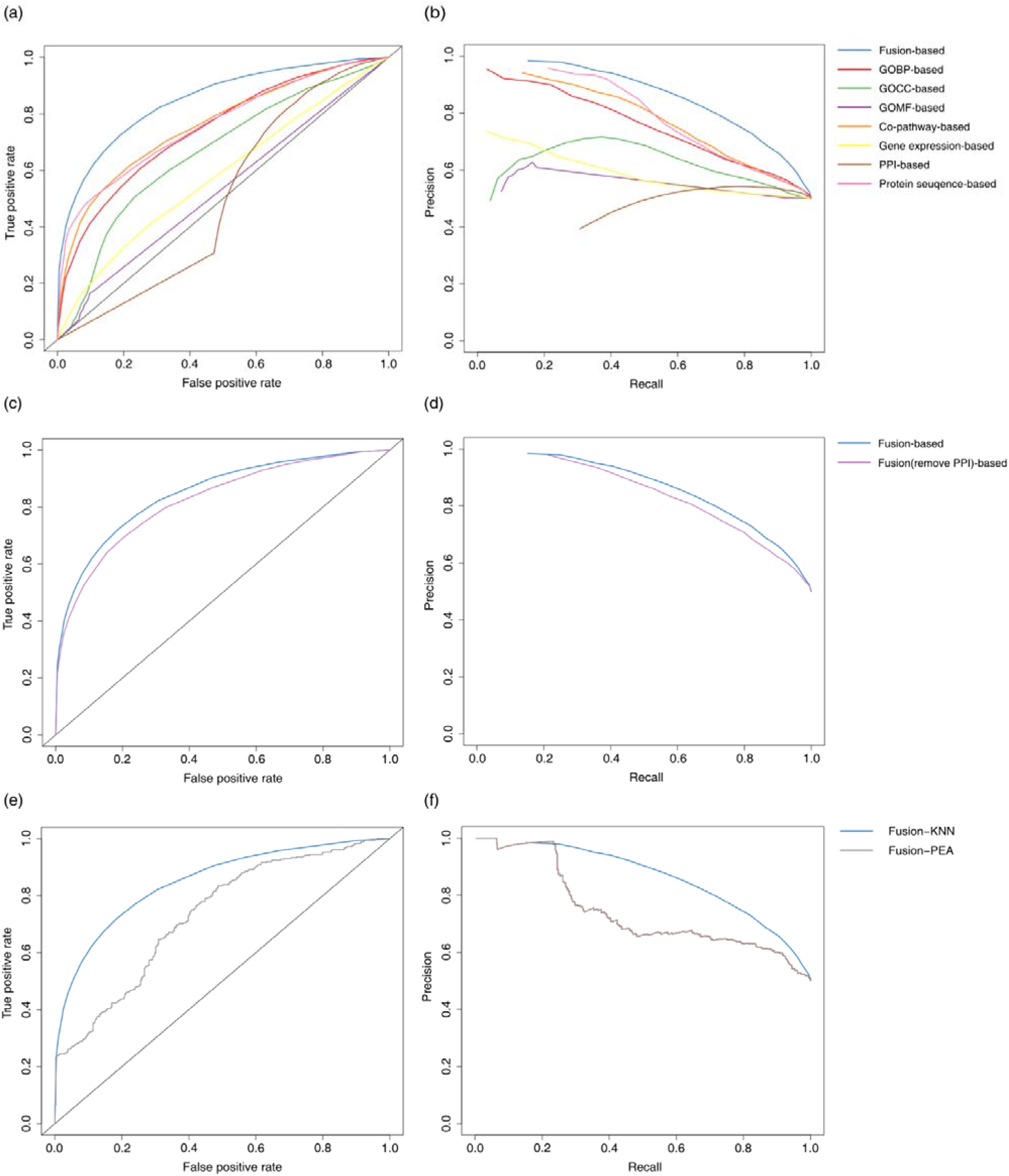
Performance of our approach. (a) ROC curve and (b) PR curve for the fusion of all seven similarity measures (blue) and the seven single similarity measures for predicting SL gene pairs. (c) ROC and (d) PR curves for the fusion of all seven similarity measures (blue) and the fusion of six similarity measures without the PPI-based (purple) for predicting SL gene pairs. (e) ROC and (f) PR curves for the k-NN algorithm (blue) and PEA algorithm (grey).

Among the model with seven single features, the sequence-based similarity measure had the best predicting performance which achieved a test AUROC of 0.76, while PPI-based similarity measure had the worst result with an AUROC of 0.50. In order to check whether PPI-based similarity measure contributes to the model, our model was further trained with the remaining six similarity measures without considering the PPI-based similarity measure. The model showed a worse performance with an AUROC of 0.82 compared to the whole-feature one (Fig.3(c)~3(d)), indicating that PPI is also a contributor to the model.

We further compared our method to the PEA algorithm based on the fusion of the seven features for their ability to predict SL gene pairs. The ROC and PR curves analysis is illustrated in Fig.3(e)~3(f), and the performance of the two methods based on the AUROC, AUPR, accuracy, precision, recall and F1 score are summarized (Table 1). According to the results, our proposed method achieves a better performance in terms of AUROC, AUPR, precision, F1 score and accuracy, which demonstrate an improvement of 13% in AUROC compared the PEA algorithm. AUROC for GOBP-based k-NN, Co-pathway-based k-NN and Protein sequence-based k-NN are 0.745, 0.761 and 0.762, respectively. AUROC for these single similarities measure-based k-NN are higher than AUROC for fusion-based PEA.

### 3.2 Comprehensive analysis and potential application of SL gene pairs

We constructed two SL networks based on the training set (Fig.4(a)) and the predicted top 3,000 SL pairs (Fig.4(b)), respectively. In the SL networks, each node represents a gene, and each edge represents a SL interaction. Node size is proportional to the number of SL pairs that the gene involved. Both in the training set (Fig.4(a)) and the predicted top 3,000 SL pairs (Fig.4(b)), we found that there are three genes with a large number of SL partners, KRAS (Entrez ID: 3845), HRAS (Entrez ID: 3265) and NRAS (Entrez ID: 4893). All the three genes are RAS genes in humans which are the most common oncogenes in human cancer. Research suggests that RAS (KRAS, NRAS and HRAS) is the most frequently mutated gene family in cancers, and mutations which permanently activate RAS are found in 20% to 25% of all human tumors, up to 90% in certain types of cancer, such as pancreatic cancer^[7]^. However, it is still a difficult task to develop an effective treatment strategy for Node size is proportional to the number of SL pairs a gene has.

**Fig.4.**
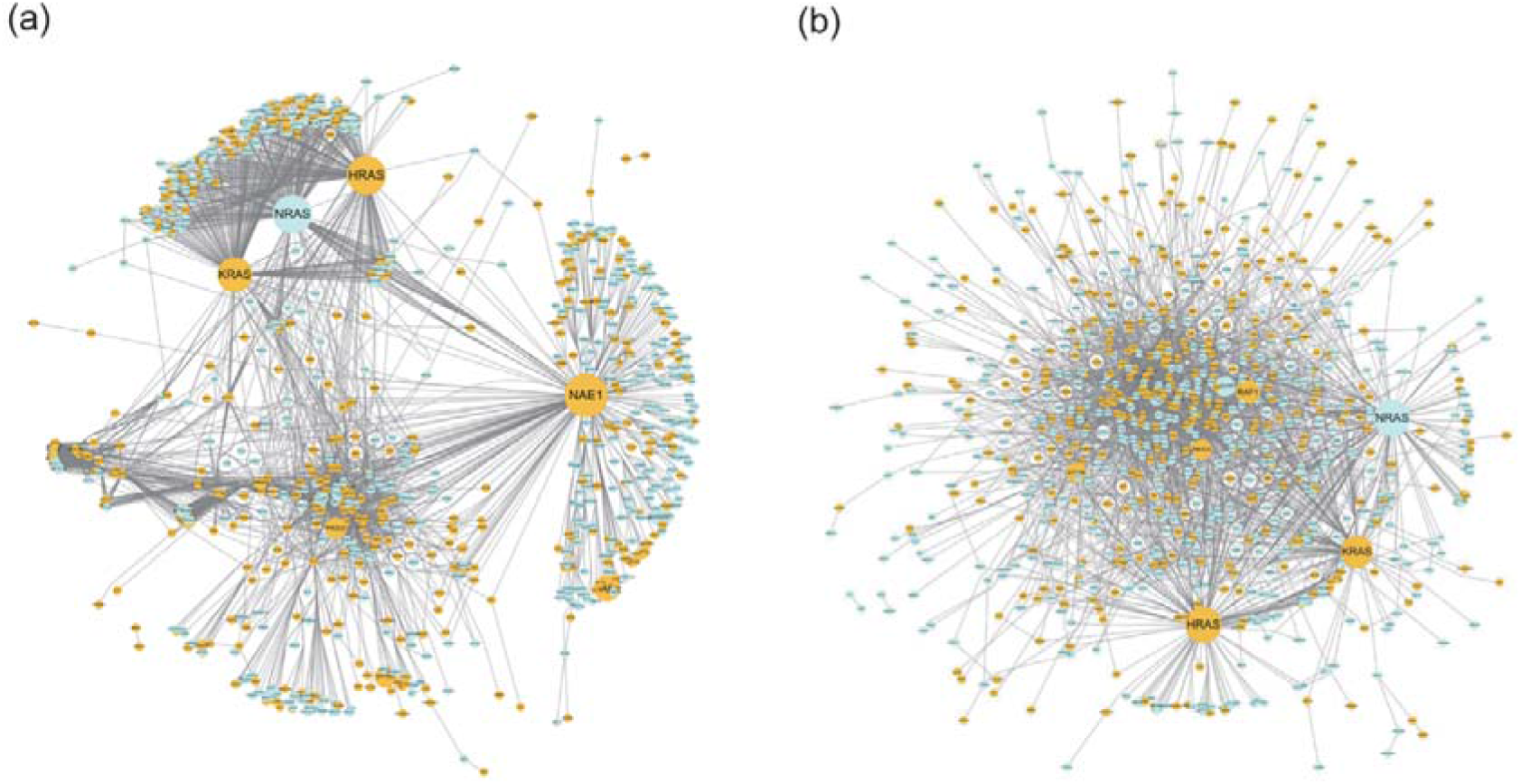
The resulting SL network in (a) training set and (b) predicted top 3,000 SL pairs. Each node represents a gene, and each edge represents a SL interaction. Yellow nodes denote druggable genes, blue nodes denote non-druggable genes.

RAS dependent cancer, the investigators have sought effective RAS inhibitors for more than three decades, but there are few available targeted drugs for RAS mutations^[39]^. For the cancer treatment, we expect that the partner genes might be promising druggable targets to achieve synthetic lethality for the cancer cells with RAS genes.

To explore the potential druggable SL partners for the RAS genes, we annotated the genes using drug target labels obtained from DrugBank^[40]^ in the SL networks (Fig.4), the druggable targets are displayed as yellow nodes. In the predicted top 3,000 SL pairs, 30 out of 251 druggable genes are the common partners of all the RAS genes (Table 2). We further employed a KEGG pathway enrichment analysis in SL partners for the RAS genes of the training set (Fig.5(a)) and the predicted top 3,000 SL pairs (Fig.5(b)). In the result, the SL partner genes of KRAS are enriched in viral infection and cancer related pathways in both training set and predicted top 3,000 SL pairs. Similar pathways are enriched for the HRAS.

**Table2.**
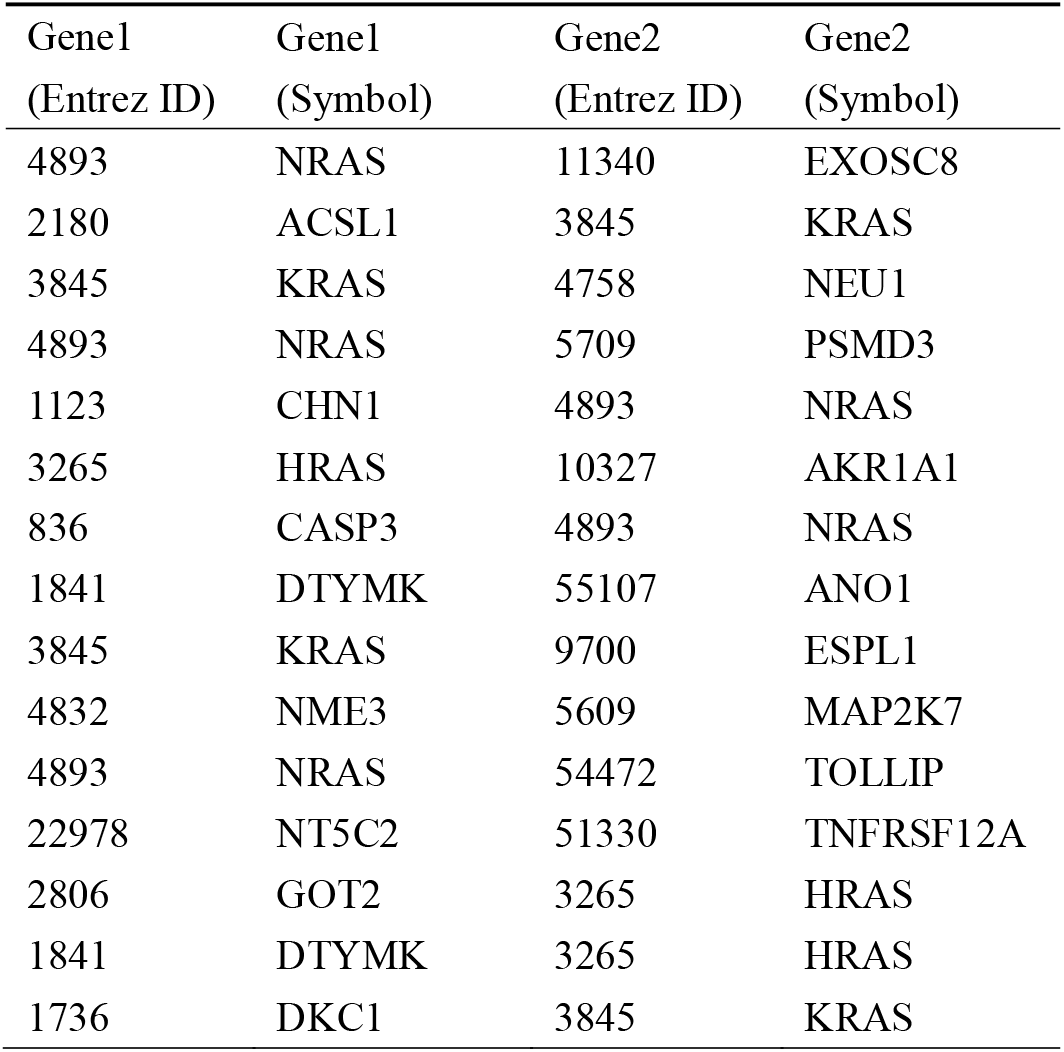
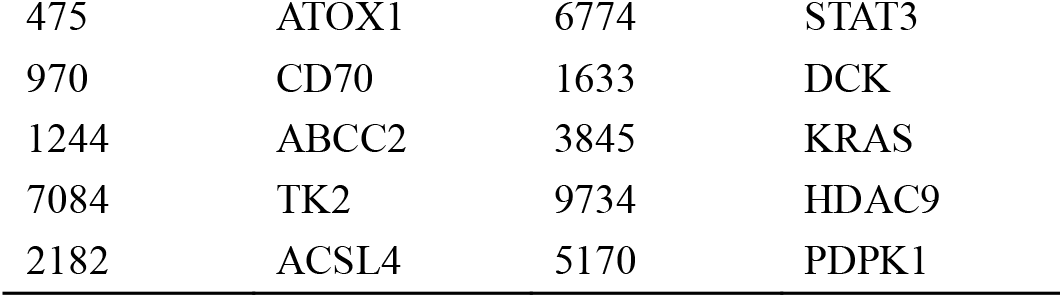
The top 20 predicted SL pairs with the higher predicted probability.

| Gene1<br>(Entrez ID) | Gene1<br>(Symbol) | Gene2<br>(Entrez ID) | Gene2<br>(Symbol) |
| --- | --- | --- | --- |
| 4893 | NRAS | 11340 | EXOSC8 |
| 2180 | ACSL1 | 3845 | KRAS |
| 3845 | KRAS | 4758 | NEU1 |
| 4893 | NRAS | 5709 | PSMD3 |
| 1123 | CHN1 | 4893 | NRAS |
| 3265 | HRAS | 10327 | AKR1A1 |
| 836 | CASP3 | 4893 | NRAS |
| 1841 | DTYMK | 55107 | ANO1 |
| 3845 | KRAS | 9700 | ESPL1 |
| 4832 | NME3 | 5609 | MAP2K7 |
| 4893 | NRAS | 54472 | TOLLIP |
| 22978 | NT5C2 | 51330 | TNFRSF12A |
| 2806 | GOT2 | 3265 | HRAS |
| 1841 | DTYMK | 3265 | HRAS |
| 1736 | DKC1 | 3845 | KRAS |
| 475 | ATOX1 | 6774 | STAT3 |
| 970 | CD70 | 1633 | DCK |
| 1244 | ABCC2 | 3845 | KRAS |
| 7084 | TK2 | 9734 | HDAC9 |
| 2182 | ACSL4 | 5170 | PDPK1 |

**Fig.5.**
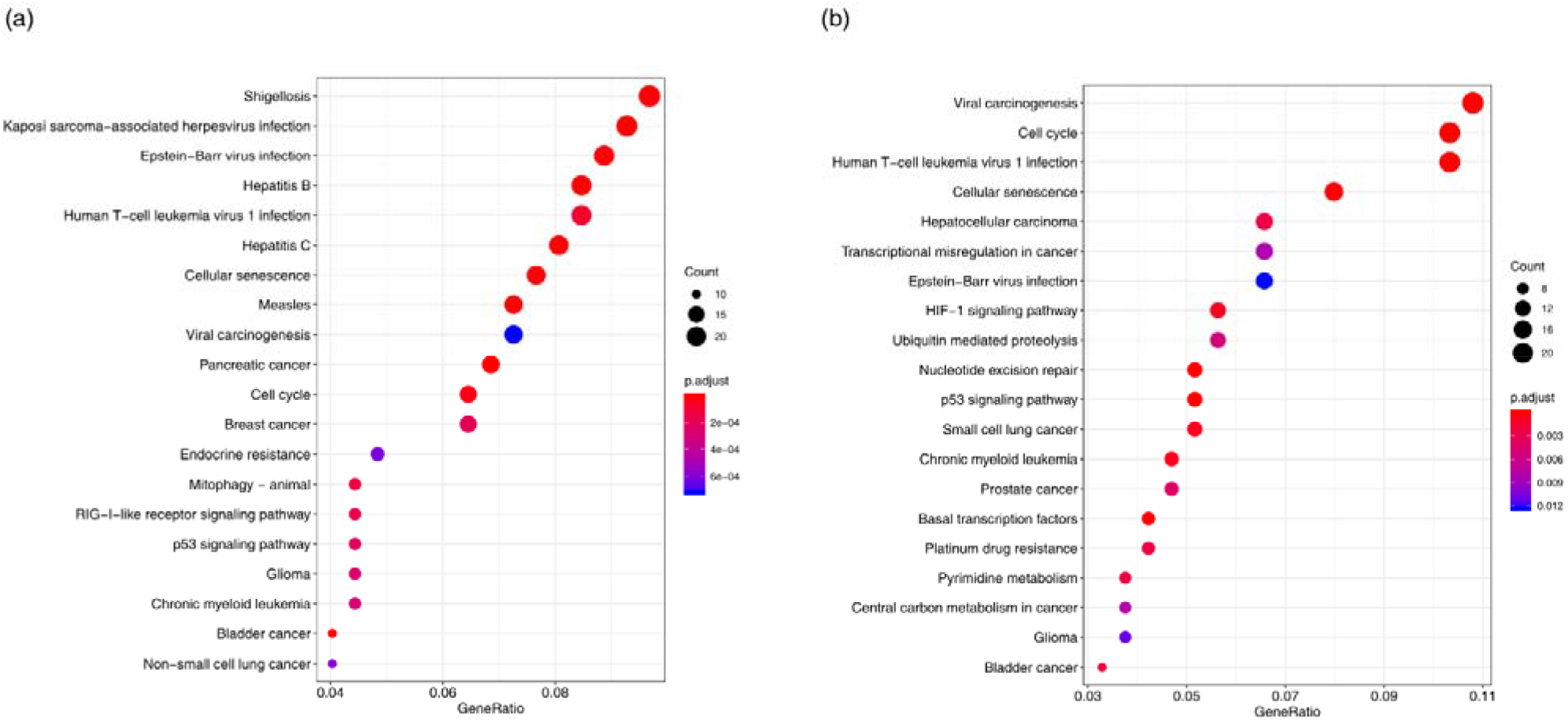
The significant enrichment KEGG pathways in (a) training set and (b) predicted top 3,000 SL pairs.

Moreover, we calculated the ATC distribution of drugs in DrugBank that can target SL partner genes for RAS genes (Fig.6). Except for the drugs without ATC code, there are more drugs with ATC code L (Antineoplastic and immunomodulating agents) than other drugs. It indicates that drugs targeting SL partner genes of RAS have been already used for cancer treatment. And RAS SL partners-targeted drugs with other ATC codes may also have anti-cancer potential. For example, Trimetrexate (PubChem ID: 5583), a drug with ATC code P (Antiparasitic products, insecticides and repellents) targets the DHFR (Entrez ID: 1719), which is an SL partner of HRAS. Researches show that Trimetrexate has potential anti-cancer activity and can be used to treat several types of cancer including colon cancer by inhibiting DHFR^[41,42]^.

**Fig.6.**
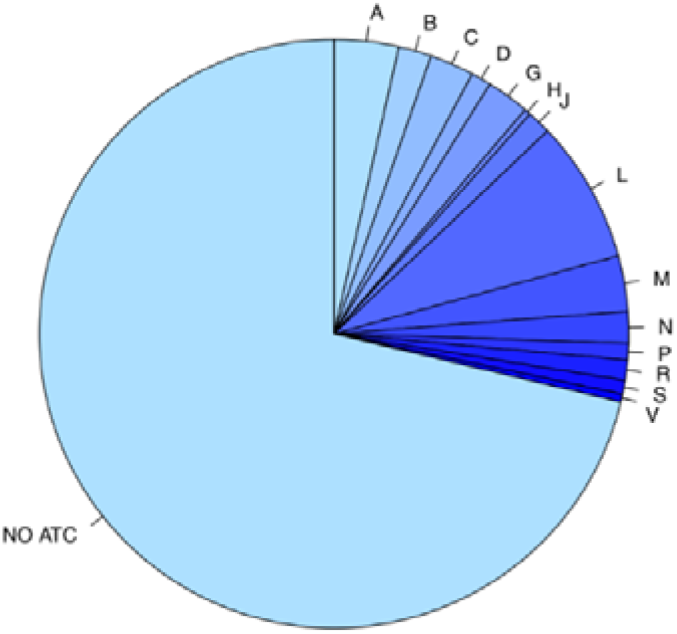
The ATC distribution of drugs in DrugBank that can target RSA-related genes’ SL partner genes.

## 4. Conclusion and Discussion

There is immense potential for synthetic lethality in cancer therapeutics. SL is considered to be the foundation of the development of the selective anticancer therapy, which aims to inhibit the SL partner of inactivated genes in cancer cells^[43,44]^. The large-scale SL screening for individual genes have been performed by RNAi and CRISPR technologies in human cell lines. But it is unfeasible for these *in vitro* screening to test the complete SL gene pairs for numerous cancers. In this work, we address such problems by developing an integrated similarity measure-based computational method to predict the SL gene pairs based on the SynLethDB database. Instead of using feature vectors directly, our approach is based on the different types of similarity measures between gene pairs, so we applied the k-NN model to achieve the prediction task. Compared to PEA, another similarity-based algorithm which get great performance in the classification task, the k-NN algorithm achieves a better performance for the prediction of SL gene pairs, which can achieve a test AUROC of 0.85.

Different from the other methods, we integrate seven types of similarity measures of the SL gene pairs into one integrated similarity measure, which greatly improves the classification performance of the model. Additionally, among these seven similarity measures, we have proved that the protein sequence- and GOBP-based similarity features showed strong predictive power, which indicate that the sequence and biological process of genes are the key element for the classification of SL pairs.

We further found that the RAS genes (i.e. KRAS, NRAS, HRAS) had the largest number of SL partners both in training set and the predicted top 3,000 SL pairs. For RAS genes, we employed a pathway enrichment analysis in SL pairs and calculated the ATC distribution of drugs that can target RSA partner genes. The results show that the RAS partner genes are enriched in pathways related to viral infection and cancer. In addition to anticancer drugs, other drugs targeting RAS SL partner genes may also have anti-cancer potential.

One limitation for this work is the limited set of samples using the intersection of sample sets with multi-dimensional features. If a certain feature of a sample is not available, the sample would be excluded from the construction of similarity network. But with the accumulation of multi-omics data in the future, the scale of sample set will be expanded and the performance of our approach will be improved. In the future work, there are more similarity measures can be added into our methods, which may further improve the performance of the model, such as SCNA data, essentiality profile data, and mutual exclusivity data. Overall, our findings suggest that our approach could be a valuable tool for predicting SL gene pairs, which may play a role in targeted therapy for cancer treatment.

